# mRNA decapping machinery targets *LBD3/ASL9* transcripts to authorize developmental reprogramming in Arabidopsis

**DOI:** 10.1101/834465

**Authors:** Zhangli Zuo, Milena Edna Roux, Jonathan Renaud Chevalier, Yasin F. Dagdas, Takafumi Yamashino, Søren Diers Højgaard, Emilie Knight, Lars Østergaard, Morten Petersen

## Abstract

Multicellular organisms perceive and transduce multiple cues to optimize developmental reprogramming and cell state switching. Key transcription factors drive developmental changes, but transitions also require the attenuation of previous states. Here, we demonstrate that the mRNA levels of the *LATERAL ORGAN BOUNDARIES DOMAIN 3 (LBD3) / ASYMMETRIC LEAVES 2-LIKE 9 (ASL9)* transcription factor are directly regulated by mRNA decapping. Capped *ASL9* transcripts accumulate in decapping deficient plants and *ASL9* mRNAs are found together with decapping components. Accumulation of *ASL9* inhibits apical hook and lateral roots formation and interestingly, exogenous auxin application restores lateral roots formation in both *ASL9* overexpressor and mRNA decay-deficient mutants. Moreover, mutations in the cytokinin transcription factors type-B ARABIDOPSIS RESPONSE REGULATORS (B-ARRs) *ARR10* and *ARR12* restore the developmental defects in apical hooking and lateral root formation caused by over-accumulation of capped *ASL9* transcript upon *ASL9* overexpression. Thus, the mRNA decay machinery directly targets *ASL9* transcripts for decay to balance cytokinin/auxin responses during developmental reprogramming.

## Introduction

Understanding normal tissue development requires information about diverse cellular mechanisms controlling gene expression. Much work has focused on of the transcriptional networks that govern stem cell differentiation. For example, ectopic expression of Yamanaka factors may lead to induced pluripotency in mice and humans[1, 2]. Similarly, ectopic expression of *LATERAL ORGAN BOUNDARIES DOMAIN (LBD)/ASYMMETRIC LEAVES2-LIKE (ASL)* genes is sufficient to induce spontaneous proliferation of pluripotent cell masses in plants, a reprogramming process triggered *in vitro* by the antagonistic phytohormones auxin and cytokinin[3, 4]. Auxin and cytokinin responses are essential for a vast number of developmental processes in plants including post-embryonic reprograming and formation of the apical hook to protect the meristem during germination in darkness[5, 6] as well as lateral root (LR) formation[7]. Loss-of-function mutants in genes that regulate auxin-dependent transcription such as *auxin-resistant1* exhibit defective hooking and LR formation[8, 9]. In addition, type-B ARABIDOPSIS RESPONSE REGULATORS (B-ARRs) ARR1, ARR10 and ARR12 work redundantly to activate cytokinin transcriptional responses in shoot development and LR formation[10–12]. Exogenous cytokinin application disrupts LR initiation by blocking pericycle founder cell transition from G2 to M phase[13]. Thus, reshaping the levels of certain transcription factors leads to changes in cellular identity. As cellular reprogramming must be tightly regulated to prevent spurious development, the expression of these transcription factors may be controlled at multiple levels[14]. However, most developmental studies focus on their transcription rates and overlook the contribution of mRNA stability or decay to these events[15].

Eukaryotic mRNAs contain stability determinants including the 5’ 7-methylguanosine triphosphate cap (m7G) and the 3’ poly-(A) tail. mRNA decay is initiated by deadenylation, followed by degradation via either 3’-5’ exosomal exonucleases and SUPPRESSOR OF VCS (SOV)/DIS3L2 or via the 5’-3’ exoribonuclease activity of the decapping complex[16, 17]. This complex includes the decapping enzymes DCP1/2 along with other factors (DCP5, DHH1, VCS, LSM1-7 complex and PAT1), and the exoribonuclease XRN that degrades monophosphorylated mRNA. mRNA decapping complex and mRNAs can aggregate into distinct cytoplasmic foci called processing bodies (PBs)[18, 19]. PAT1 (Protein Associated with Topoisomerase II, PAT1b in mammals) promotes PB assembly and activates decapping by binding mRNA and recruiting other decapping components[20–22].

mRNA decay regulates mRNA levels and thereby impacts cellular reprogramming[23, 24]. We and others have shown that the decapping machinery is involved in stress and immune responses[25–30], and that RNA binding proteins can target selected mRNAs for decay[29–31]. Postembryonic lethality[32] and stunted growth phenotypes[33, 34] associated with disturbance of the decay machinery indicate the importance of mRNA decapping and decay machinery during plant development. However, while much has been learned about how mRNA decapping regulates cellular reprogramming during plant stress responses[29, 30], far less is known about how decapping contributes to plant development.

While developmental defects and altered transcriptomes are well described, *dcp1*, *dcp2* and *vcs* mutants display postembryonic lethality whereas *lsm1a/lsm1b* and *dcp5* mutants exhibit abnormal development including cotyledons with disorganized veins while *lsm1a/lsm1b* are dwarfs and *dcp5* displays a delayed growth phenotype[32–34]. All these differences suggest that mutations in mRNA decay components may cause pleiotropic phenotypes not directly linked to mRNA decapping and decay deficiencies [26, 35, 36]. For example, it has been proposed that lethality in some mRNA decay loss-of-function mutants is not due to decay deficiencies *per se*, but to the activation of immune receptors which evolved to surveil microbial manipulation of the decay machinery[26]. In line with this, loss-of-function of *AtPAT1* inappropriately triggers the immune receptor SUMM2, and *Atpat1* mutants consequently exhibit dwarfism and autoimmunity[26]. Thus, PAT1 is under immune surveillance and PAT1 function(s) are best studied in SUMM2 loss-of-function backgrounds.

Here we disrupt the mRNA decapping components PAT1 and its 2 paralogues PATH1 and PATH2 to study the impact of mRNA decay during developmental reprogramming. By disrupting the decapping machinery in the *summ2* background[26] we can study this process without autoimmunity disturbance. Our approach, together with analyses of the *dcp2* and *dcp5* mRNA decay-deficient mutants, reveals that the mRNA decay machinery directly regulates the important developmental regulator *ASL9*. Thus, when mRNA decay is disrupted, *ASL9* accumulates and inhibits cells from forming apical hooks and lateral roots. Moreover, interference with a cytokinin pathway and/or exogenous auxin application restores the developmental defects in both *ASL9* over-expressing plants and in mRNA decay deficient mutants. These observations indicate that the mRNA decay machinery is fundamental to cellular reprogramming during developmental decision making.

## Results

### PAT mRNA decay factors are needed to initiate developmental processes

We previously showed that the PAT1 decapping factor localizes in PBs and that the immune receptor SUMM2 is triggered in the absence of PAT1. In addition to PAT1, two other PATs, PATH1 and PATH2, are encoded by the *Arabidopsis* genome[26]. We therefore initially confirmed that the PAT1 paralogues PATH1 and PATH2 also localize to PBs and interact with the LSM1 decapping factor (Fig S1A). To examine if impaired mRNA decay stalls developmental reprogramming, we developed a triple knockout of *PAT1*, *PATH1* and *PATH2* in the *summ2* background (Fig 1A, S1B-F). This permits analyses of decapping deficiency in an autoimmune free background [26] (Fig S1G). The growth phenotypes of 6 weeks-old soil-grown plants of *pat* single, double and triple mutants at 21 °C with 8 hrs photoperiod are shown in Fig. 1A&S1B. *path1/summ2, path2/summ2* and *path1/path2/summ2* have leaves similar to *summ2* while *pat1/summ2, pat1/path1/summ2, pat1/path2/summ2* and *pat1/path1/path2/summ2* exhibit serrated leaves. Compared to *pat1/summ2*, leaf serration and dwarfism are more severe in *pat1/path1/summ2,* while *pat1/path2/summ2* exhibits less serration but longer petioles. *pat1/path1/path2/summ2* mutants exhibited markedly stunted growth compared to the other *pat* single or double mutants (Fig 1A, S1B-F). To identify genes which affect different developmental programs regulated by mRNA decapping, we performed RNA-seq from plants of *pat1/path1/path2/summ2* and all single and double mutant combinations. Data File S1 and Fig 2B exhibit differently expressed genes in *pat1/path1/path2/summ2* which are clustered in Fig 1C and annotated in Fig 1D&E [37]. This cluster analysis of the mRNA decapping deficient mutants showed that *i*. genes involved in oxidation-reduction are largely misregulated, while *ii*. transcripts involved in oxidative stress response accumulated, and *iii*. transcripts responsible for auxin response and signaling, transcription regulation and growth were reduced (Fig 1D&E).

**Fig 1.**
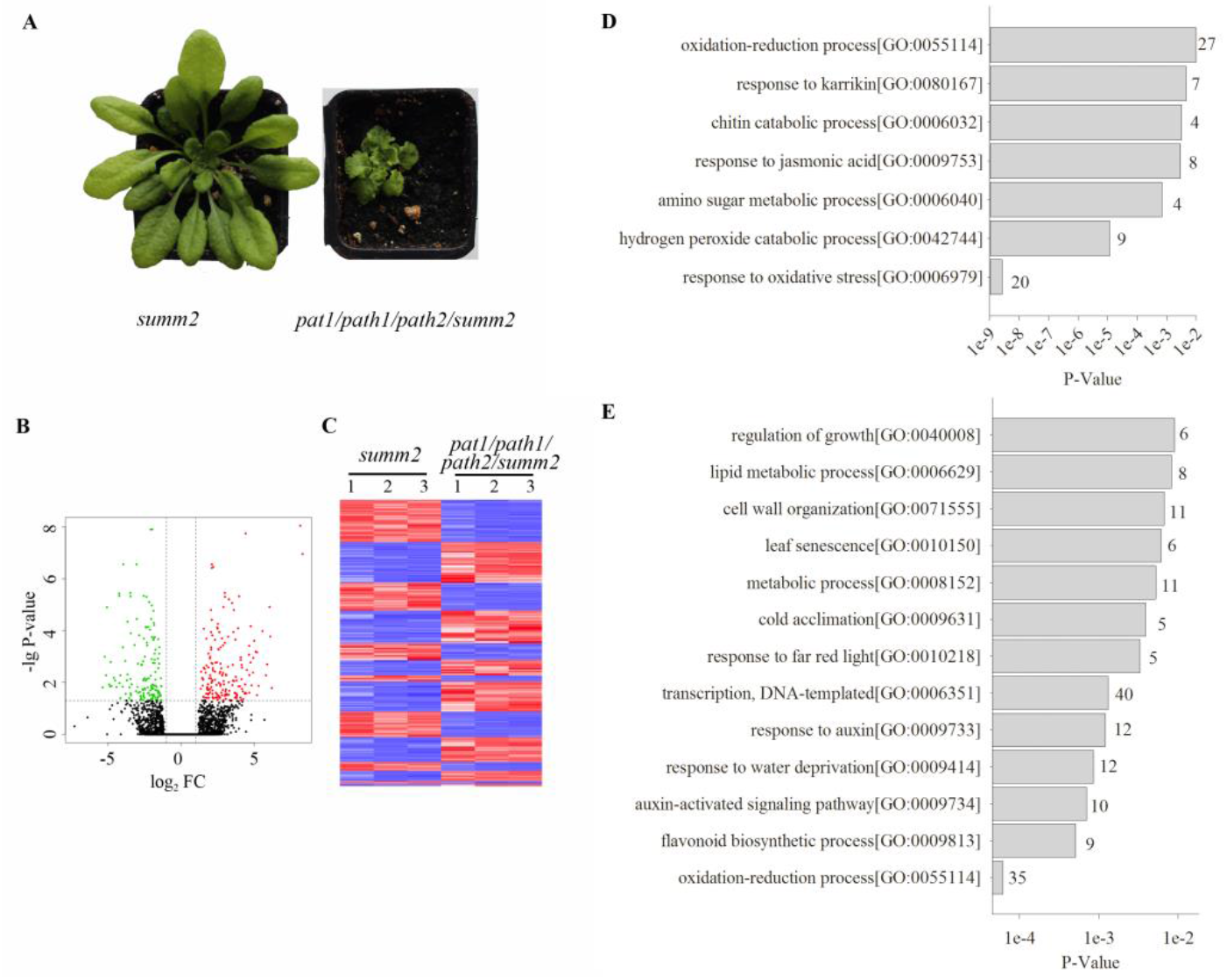
mRNA decay component PATs are needed for transcript removal and developmental processes initiation. (**A**) 6 week-old plants of *summ2* and *pat1/path1/path2/summ2* grown in soil in a chamber with 8/16hrs light/dark at 21°C. A representative plant for each line is shown and pictures in Fig S1B were taken at the same time. (B) Volcano plot for differently expressed genes in *pat1/path1/path2/summ2* compare to *summ2*. Up-regulated genes (FC≥2 (log2(FC)≥1), P-value≤0.05) are coloured red and down-regulated (FC≤2 (log2(FC)≤-1), P-value≤0.05) green. (C) Heat map clustering of differentially expressed genes for comparison of *pat1/path1/path2/summ2* and *summ2*. Pearson’s metrics were used in hierarchical clustering of the genes. In the plot, red indicates high expression and blue low expression. Gene ontology of transcripts upregulated (**D**) or downregulated (**E**) in 6 week-old plants of mRNA decay deficient mutant *pat1/path1/path2/summ2* compared to *summ2*.

**Fig 2.**
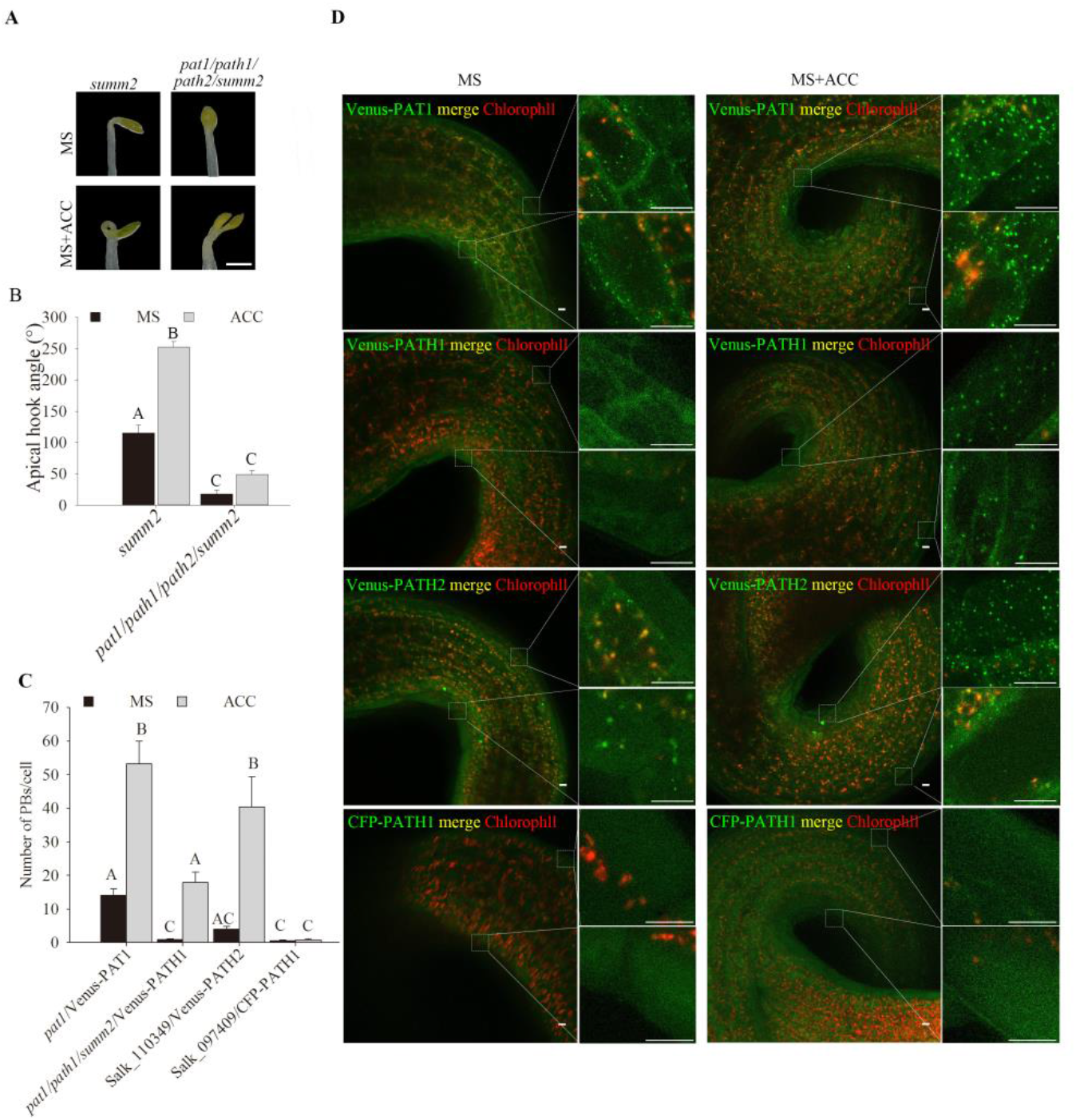
*PAT* loss-of-function causes deregulation of cell decision making during apical hooking. Hook phenotypes (**A**) and apical hook angles (**B**) in triple responses to ACC treatment of etiolated *summ2* and *pat1/path1/path2/summ2* seedlings. The experiment was repeated 3 times, and representative pictures are shown. The scale bar indicates 1mm. (**C**) Quantification of PBs/cell and (D) representative confocal microscopy pictures of hook regions following ACC treatment. 4 day-old dark-grown *pat1*/Venus-PAT1, *pat1/path1/summ2/*Venus-PATH1, Salk_110349/Venus-PATH2 and Salk_097409/CFP-PATH1, seedlings on MS or MS+ACC plates for 4 days. Scale bars indicate 10 μm. Bars marked with the same letter are not significantly different from each other (P-value>0.05).

### *PAT* loss-of-function causes deregulation of apical hooking

Our RNA-seq analysis indicates mRNA decay is needed to remove superfluous transcripts which may affect cellular decision making and developmental reprogramming. To assess this, we explored readily scorable phenotypic evidence of defective decision making during development. Since apical hooking can be exaggeratedly induced by exogenous application of ethylene or its precursor ACC, we germinated seedlings in darkness in the presence or absence of ACC [38, 39]. Interestingly, of all lines tested only *pat1/path1/path2/summ2* was hookless and unable to make the exaggerated apical hook under ACC treatment (Fig 2A, B&S2A). This indicates that mRNA decapping is required for the commitment to apical hooking. Supporting this notion, ACC treatment led to a massive increase of Venus-PAT1 and Venus-PATH2 foci in hook regions in their corresponding *pat1* and *path2* mutants (Salk_110349) backgrounds. In contrast, CFP-PATH1 exhibited no change in distinct foci number (Fig2C&D), although Venus-PATH1 foci increased in the hook region in ACC treated *pat1/path1/summ2* triple mutants. Collectively, these data show that PATs may be involved in ethylene-induced apical hook formation.

### PAT decay components target *ASL9*

To search for transcripts responsible for the hookless phenotype, we revisited our RNA-seq data (Data Set S1) and verified that transcripts of *RSM4 (RADIALIS-LIKE SANT/MYB)* and of *ASL9* (*ASYMMETRIC LEAVES 2-LIKE 9*, also named *LBD3, LOB DOMAIN-CONTAINING PROTEIN 3*) accumulated specifically in *pat1/path1/path2/summ2* mutants (Data Set S1). Overexpression of the close *RSM4* homologue *RSM1* prevents apical hook formation [40]. However, qPCR analysis of *RSM4* transcripts revealed that it accumulated in several *pat* mutants which were able to develop apical hooks (Fig S2C). This indicates that over-accumulation of *RSM4* does not contribute to the hookless phenotype of *pat1/path1/path2/summ2*.

ASL9 belongs to the large AS2/LOB (ASYMMETRIC LEAVES 2/LATERAL ORGAN BOUNDARIES) family [41] which includes key regulators of organ development [42]. Interestingly, the ASL9 homologue ASL4 negatively regulates brassinosteroid accumulation to limit growth in organ boundaries, and overexpression of *ASL4* impairs apical hook formation and leads to dwarfed growth [43]. While *ASL4* mRNA did not accumulate in *pat1/path1/path2/summ2* mutants, we hypothesized that ASL9 could also interfere with hook formation. We therefore analyzed mRNA levels of *ASL9* in ACC-treated seedlings and verified that *pat1/path1/path2/summ2* accumulated at least 2-fold higher levels of *ASL9* transcript compared to any other lines (Fig 3A, S2D). Concordantly, an over-expressor line (Fig 3B) of *ASL9* (Col-0/*oxASL9*) [44] also exhibited a hookless phenotype (Fig 3C&D). However, we did not observe any changes including tighter apical hooks in *asl9-1 (SAIL_659_D08)* mutants (Fig S2E-G) suggesting other members of the AS2/LOB family act redundantly in this process. Nevertheless these results indicate that apical hook formation in *pat1/path1/path2/summ2* is compromised due to misregulation of *ASL9*.

**Fig 3.**
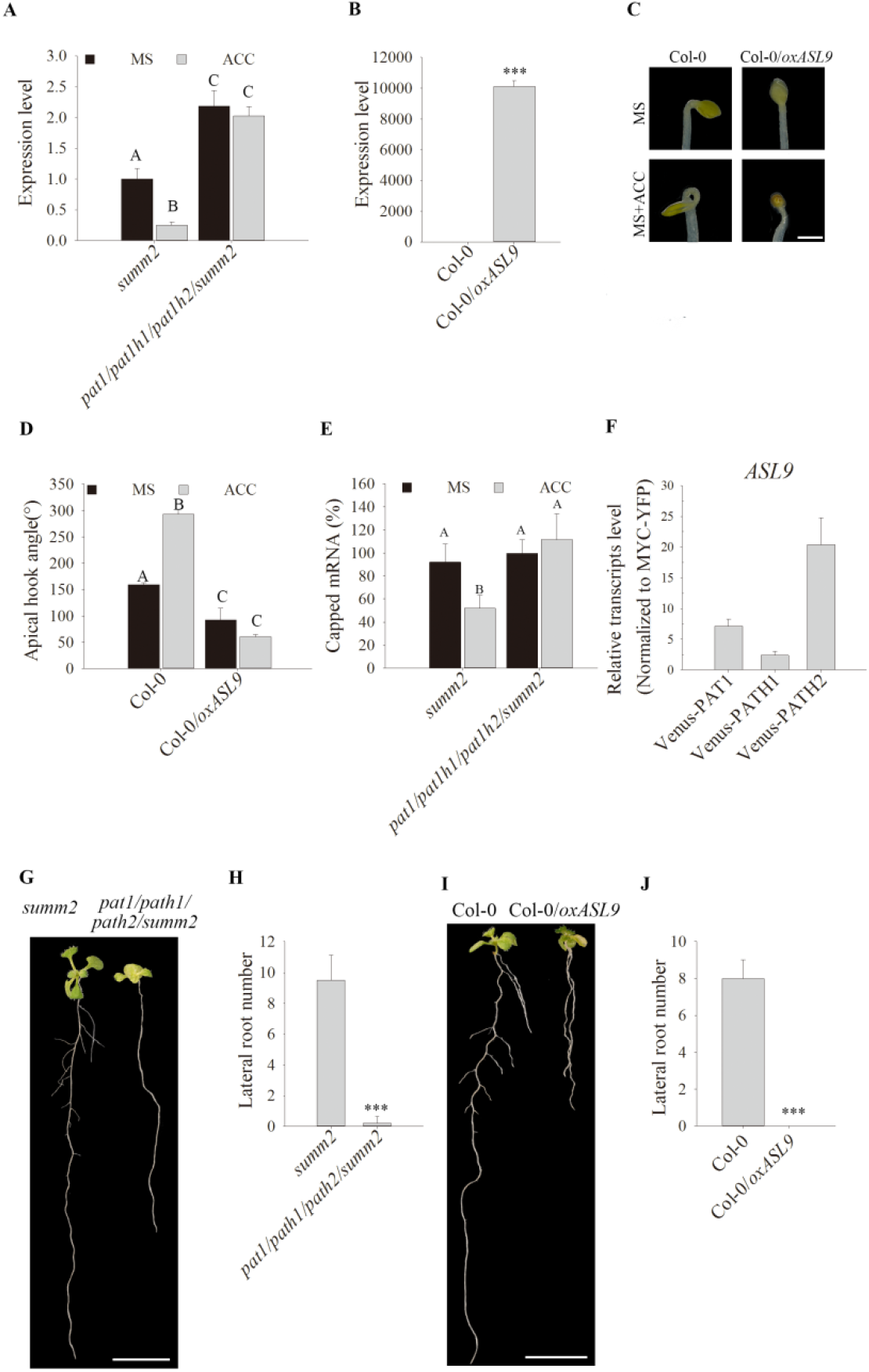
PATs target *ASL9* to regulate apical hooking and LR formation. (**A**) *ASL9* mRNA levels in cotyledons and hook regions of dark-grown *summ2* and *pat1/path1/path2/summ2* seedlings under control or ACC treatment. Error bars indicate SE (n = 3). One representative of 3 biology replicates is shown. (**B**) *ASL9* expression levels in 4 day-old seedlings of Col-0 and Col-0/*oxASL9*. Error bars indicate SE (n = 3). One representative of 3 biology replicates is shown. Hook phenotypes (**C**) and apical hook angles (**D**) of triple response to ACC treatment of etiolated seedlings of Col-0 and Col-0/*oxASL9*. The experiment was repeated 3 times, and representative pictures are shown. The scale bar indicates 1mm. (**E**) Capped *ASL9* transcript levels in cotyledons and hook regions of dark-grown *summ2* and *pat1/path1/path2/summ2* seedlings. Error bars indicate SE (n = 3). (**F**) All 3 PATs can bind *ASL9* transcripts. 4-day dark-grown plate seedlings of *pat1*/Venus-PAT1, *pat1/path1/summ2*/Venus-PATH1 and *pat1/path2/summ2*/Venus-PATH2 were taken for the RIP assay. *ASL9* transcript levels were normalized to those in RIP of MYC-YFP as a non-binding control. Error bars indicate SE (n=3). Phenotypes (**G**) and LR number counts (**H**) of 10-day old seedlings of *summ2* and *pat1/path1/path2/summ2*. The experiment was repeated 4 times, and representative pictures are shown. The scale bar indicates 1cm. Pictures in Fig S4A were taken at the same time. Phenotypes (**I**) and LR number counts (**J**) of 10-day old seedlings of Col-0 and Col-0/*oxASL9*. Treatment was repeated 3 times, and representative pictures are shown. The scale bar indicates 1cm. Bars marked with three asterisk (***) are statistically extremly significant (P-value <0.001) and bars marked with the same letter are not significantly different from each other (P-value>0.05).

To determine whether *ASL9* is a target of the decapping complex, we assayed for capped *ASL9* transcripts in ACC and mock-treated *pat* mutants. By calculating the ratio between capped versus total *ASL9* transcripts, we verified that with ACC treatment, *pat1/path1/path2/summ2* accumulated significantly higher levels of capped *ASL9* transcripts than all lines but *pat1/path2/summ2* (Fig 3E, S2D), although only the former was statistically different from the *summ2* control. Moreover, RNA immunoprecipitation (RIP) revealed enrichment of *ASL9* in all Venus-PAT lines compared to a free YFP control line (YFP-WAVE) (Fig 3F), being highest for PATH2 and lowest for PATH1. These data confirm that *ASL9* mRNA can be found in PAT complexes, and that mRNA decapping contributes to ACC-induced apical hook formation by regulating *ASL9* mRNA levels, preferentially but not exclusively by PAT1 and PATH2.

### Accumulation of *ASL9* suppresses LR formation

Lateral root (LR) formation is another example of post embryonic decision making. In *Arabidopsis* the first stage of LR formation requires that xylem pole cells change fate to become LR founder cells, a process positively regulated by auxin and negatively regulated by cytokinin and ethylene [45, 46]. We therefore examined LR formation in *pat* mutants and in Col-0/*oxASL9*, as *ASL9* has been implicated in cytokinin signaling [44] and as the auxin efflux gene *PIN5*, auxin induced gene *SAUR23* and *IAA19* and auxin biosynthesis gene *TAR2* are repressed in both *pat1/path1/path2/summ2* and Col-0/*oxASL9* (Fig S3). While LR formation was reduced in *pat1/path1/summ2* and *pat1/path2/summ2*, it was almost absent in *pat1/path1/path2/summ2* and *ox/ASL9* (Fig 3G-J, S4A&B). However, similar to normal hook development lateral root formation in *asl9-1* mutants appeared normal (Fig S4C&D). Interestingly, upon close inspection of *pat1/path1/path2/summ2* quadruple mutant roots (n=5), we did not observe any asymmetric cell divisions of the pericycle or indeed LR primordia in later stages[47].

### Core mRNA decapping components are involved in regulating *ASL9* transcripts

PAT mRNA decay components are needed to unlock apical hooking and LR formation, indicating the mRNA decay machinery is involved in these processes. We therefore examined apical hook and LR formation in 2 other mRNA decay deficient mutants, *dcp2* [32] and *dcp5 [33]*. Etiolated 4-day old *dcp2* and *dcp5* seedlings also exhibited a hookless phenotype (Fig 4A&B). Since *dcp2* mutants are postembryonic lethal, we used seeds from a parental heterozygote but genotyping showed all hookless seedlings were *dcp2* homozygotes (Fig S5A). Similar to observations in *pat1/path1/path2/summ2*, *ASL9* expression is not affected by ACC treatment of *dcp2* and *dcp5* seedlings while expression is lower in ACC treated Col-0 seedlings compared to untreated controls (Fig 4C). In line with this, capped *ASL9* transcripts also accumulate in ACC treated *dcp2* and *dcp5* seedlings (Fig 4D) and LR formation is also significantly reduced in *dcp5* mutants (Fig 4E& F). Thus, regulation of *ASL9* transcript levels involves core components of the mRNA decapping machinery.

**Fig 4.**
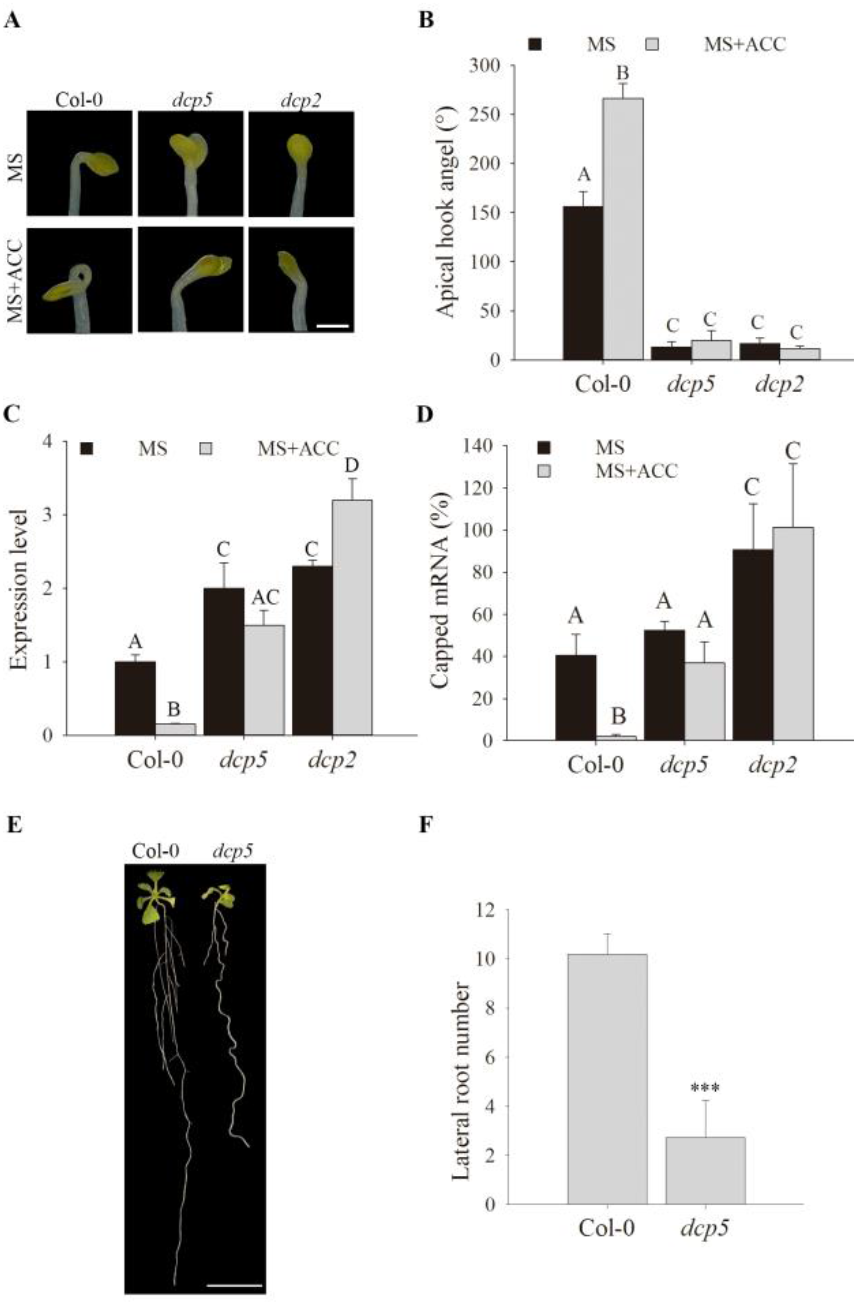
*ASL9* transcripts accumulate in many mRNA decay mutants with defects in apical hook and LR formation. Hook phenotypes (**A**) and apical hook angles (**B**) in triple responses to ACC treatment of etiolated Col-0, *dcp5* and *dcp2* seedlings. The treatment was repeated 3 times, and representative pictures are shown. The scale bar indicates 1mm. (**C**) *ASL9* mRNA levels in cotyledons and hook regions of dark-grown Col-0, *dcp5* and *dcp2* seedlings under control or ACC treatment. Error bars indicate SE (n = 3). One representative of 3 biology replicates is shown. (**D**) Capped *ASL9* transcript levels in cotyledons and hook regions of dark-grown Col-0, *dcp5* and *dcp2*seedlings. Error bars indicate SE (n = 3). Phenotypes (**E**) and LR number counts (**F**) of 10-day old seedlings of Col-0 and *dcp5*. Treatment was repeated 3 times, and representative pictures are shown. The scale bar indicates 1cm. Bars marked with three asterisk (***) are statistically extremly significant (P-value <0.001) and bars marked with the same letter are not significantly different from each other (P-value>0.05).

### Interference of a cytokinin pathway and/or exogenous auxin restores developmental defects of *ASL9* over-expressor and mRNA decay deficient mutants

*ASL9* has been implicated in cytokinin signaling [44] in which ARR1, ARR10 and ARR12 are responsible for activation of cytokinin transcriptional responses [10, 11] and cytokinin acts antagonistically with auxin. Apical hooking and lateral root formation are classic examples of auxin dependent developmental reprogramming [48]. Since auxin responsive genes are repressed in both mRNA decay mutants and in Col-0/*oxASL9*, we hypothesized that the developmental defects of mRNA decay mutants and Col-0/*oxASL9* are due to repressed auxin responses possibly caused by strong cytokinin signaling. To test this, we examined the developmental phenotype of *ASL9* over-expressors in *arr10*/*arr12* mutants [10]. Interestingly, both apical hooking and lateral root formation phenotypes were largely restored in this background (Fig 5&S5B). We also applied exogenous auxin to mRNA decay mutants *pat1/path1/path2/summ2* and *dcp5* and Col-0/*oxASL9*. This showed that 0.2 μM IAA could partially restore LR formation in *pat1/path1/path2/summ2* and *dcp5* while 2 μM IAA could partially restore LR formation of Col-0/*oxASL9* (Fig S6). These findings indicate that the mRNA decay machinery targets *ASL9* to keep cytokinin/auxin responses balanced during development.

**Fig 5.**
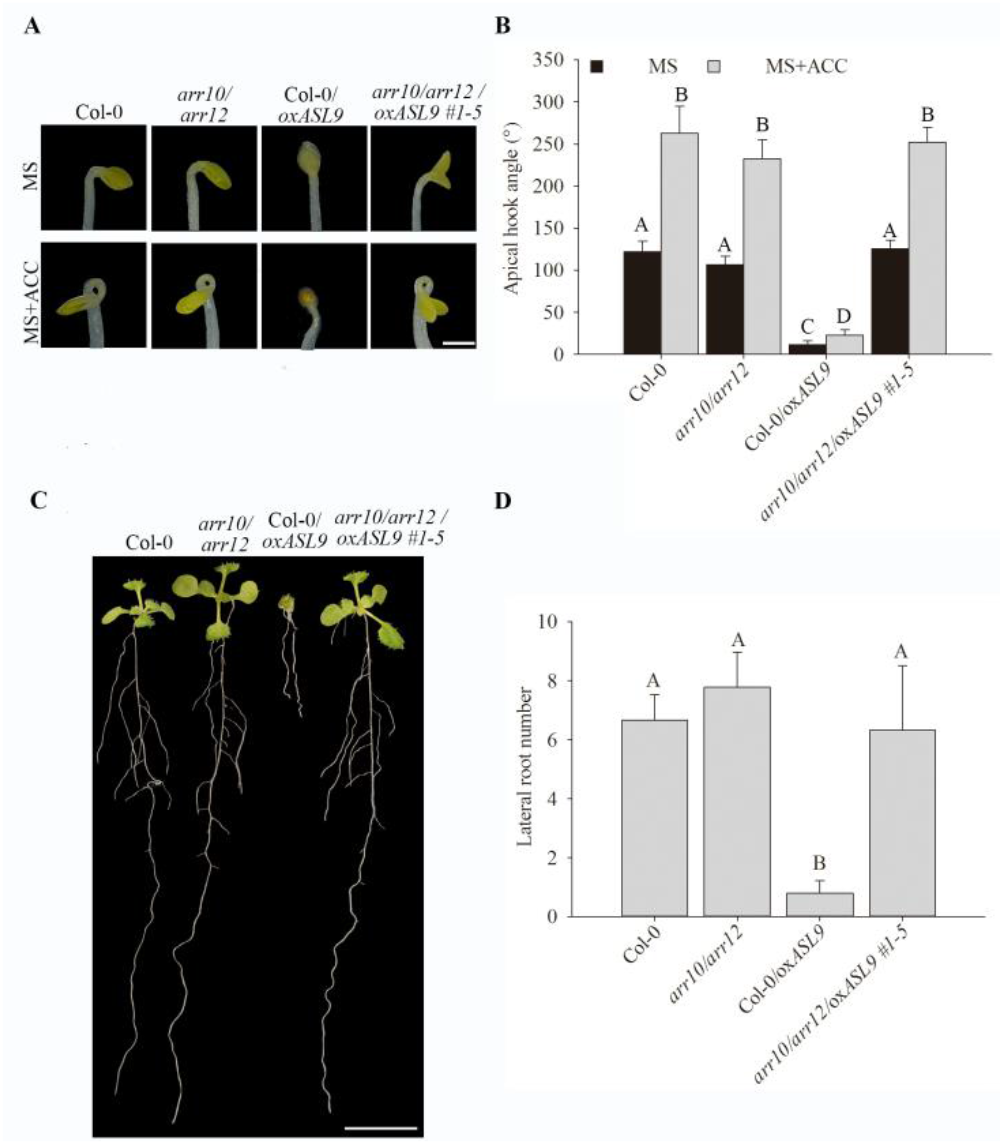
*ARR10* and *ARR12* loss-of-function restores apical hook and LR formation in *ASL9* over-expressor plants. Hook phenotypes (**A**) and apical hook angles (**B**) in triple responses to ACC treatment of etiolated Col-0, *arr10/arr12,* Col-0*/oxASL9* and *arr10/arr12/oxASL9* seedlings. The treatment was repeated 3 times, and representative pictures are shown. The scale bar indicates 1mm. Phenotypes (**C**) and LR number counts (**D**) of 10-day old seedlings of Col-0 *arr10/arr12,* Col-0*/oxASL9* and *arr10/arr12/oxASL9*. Treatment was repeated 3 times, and representative pictures are shown. The scale bar indicates 1cm. Bars marked with the same letter are not significantly different from each other (P-value>0.05).

## Discussion

Cellular reprograming requires massive overhauls of gene expression [49]. Apart from unlocking effectors needed to install a new program, previous states or programs also need to be terminated [14]. We report here that mRNA decay is required to unlock cellular states during development. The stunted growth phenotype and downregulation of developmental and auxin responsive mRNAs in the *pat1/path1/path2/summ2* mutant (Fig 1C) supports a model in which defective clearance of suppressors of development hampers decision making upon hormonal perception. Apical hooking and LR formation are classic examples of auxin-dependent developmental reprogramming [48]. In line with this, we and others observed that mRNA decay-deficient mutants are impaired in apical hooking (Fig 2A, B, 4A&B) and LR formation (Fig 3G, H, 4E&F) [34, 50]. Interestingly, among the transcripts upregulated in these decay-deficient mutants was that of capped *ASL9/LBD3* (Fig 3A&4C), which is involved in cytokinin signaling [44]. Cytokinin and auxin act antagonistically [51], and cytokinin can both attenuate apical hooking [52] and directly affect LR founder cells to prevent initiation of lateral root primordia [53]. Our findings that defective reprogramming during those developmental events in mRNA decay-deficient mutants involves *ASL9* was supported by our observation that *ASL9* mRNA is directly regulated by the decapping machinery and that *oxASL9* transgenic lines cannot reprogram to attain an apical hook or to form LRs (Fig 3). In line with this, we speculate that the inability to terminate cytokinin-dependent programs prevents the correct execution of auxin-dependent reprograming in mRNA decay-deficient mutants. This is supported by the observation that treating *pat1/path1/path2/summ2*, *dcp5* and Col-0/*oxASL9* with exogenous auxin partially restores LR formation (Fig S6), and that the defects in both apical hooking and LR formation of *ASL9* over-expressing plants are largely restored by internally knocking out the cytokinin signaling activator genes *ARR10* and *ARR12* (Fig 5).

Arabidopsis contains 42 *LBD/ASL* genes[41], among these genes *LBD16*, *LBD17*, *LBD18* and *LBD29* control lateral roots formation and regulate plant regeneration[3] and overexpression of another member *ASL4* also impairs apical hook[43]. Constitutive *ASL9* expression is sufficient to suppress apical hook and LR formation, however we did not observe tighter apical hook and more lateral roots in *asl9-1* which is most likely due to functional redundancy within the *LBD/ASL* family. The accumulation of capped *ASL9* transcripts in *pat* triple mutants after ACC treatment indicates that all three PATs target *ASL9* for decay. Our observation that all three PATs can re-localize to PBs in the hook region during the triple response supports this model. Intriguingly, the detection in ACC-treated hook region PBs of Venus-PATH1 in the hook region of ACC-treated *pat1/path1* mutant seedlings only may exemplify the compensatory activities of subfunctionalized paralogues [54, 55].

Deadenylated mRNA can be degraded via either 3’-5’ exosomal exonucleases and SUPPRESSOR OF VCS (SOV)/DIS3L2 or via the 5’-3’ exoribonuclease activity of the decapping complex [16, 17]. Sorenson et al. (2018) found that *ASL9* expression is dependent on both VCS and SOV based on their transcriptome analysis, so that *ASL9* might be a substrate of both pathways [17]. While more direct data is needed to conclude whether SOV can directly regulate *ASL9* mRNA levels, we have shown that *ASL9* is a target of the mRNA decapping machinery. The function of PBs in mRNA regulation has been controversial since mRNAs in PBs may be sequestered for degradation or re-enter polysomal translation complexes [56]. A recent study has confirmed that PBs function as mRNA reservoirs in dark-grown seedlings [50]. However, our finding of direct interaction of *ASL9* transcripts and the PATs, together with the accumulation of capped *ASL9* in mRNA decay mutants, indicates that *ASL9* misregulation in *pat1/path1/path2/summ2*, *dcp2* and *dcp5* mutants is due to mRNA decapping deficiency (Fig 3A, E, F&4A-D).

## Materials and Methods

### Plant materials and growth conditions

*Arabidopsis thaliana* ecotype Columbia (Col-0) was used as a control. T-DNA insertion lines for At1g79090 (PAT1) *pat1* (Salk_040660), At1g12280 (SUMM2) *summ2-8* (SAIL_1152A06) and double mutant of PAT1 and SUMM2 *pat1/summ2* have been described [26, 57, 58], T-DNA insertion lines for AT5G13570(DCP2) *dcp2* (Salk_000519), At1g26110 (DCP5) *dcp5* (Salk_008881) and double mutant *arr10/arr12* has also been described [10, 32, 33]. The T-DNA line for PATH1 (AT3g22270) is Salk_097409 with an insertion in the 5’-UTR,for PATH2 (AT4g14990) is Salk_110349 with an insertion in the last exon and for ASL9 (AT1g16530) is SAIL_659_D08 with insertion in the first exon. Primers for newly described T-DNA lines are provided in Table S1. *pat1/path1/summ2* and *path2/summ2* mutants were generated using CRISPR/CAS9 following standard procedures with plasmid pHEE401 containing an egg cell-specific promoter to control CRISPR/CAS9 [59]. Cas9-free T2 plants were used in crosses to produce *path1/summ2, pat1/path2/summ2, path1/path2/summ2, pat1/path1/path2/summ2* mutants. Homozygous F2 plants were used for experiments. The YFP WAVE line was from NASC (Nottingham, UK) [60]. Col-0/*oxASL9* line has been described before [44].

Plants were grown in 9×9cm or 4×5cm pots at 21°C with 8/16hrs light/dark regime, or on plates containing Murashige–Skoog (MS) salts medium (Duchefa), 1% sucrose and 1% agar with 16/8hr light/dark.

### Plant treatments

For ethylene triple response assays, seeds were plated on normal MS and MS+50μM ACC, vernalized 96hrs and placed in the dark at 21°C for 4 days before pictures were taken. Cotyledon and hook regions of etiolated seedlings were collected after placing in the dark at 21°C for 4 days for gene expression and XRN1 assay. For LR formation assays, seeds on MS plates were vernalized 96hrs and grown with 16/8 hrs light/dark at 21°C vertically for 10 days. For external IAA application for LR formation experiments, seeds on MS plates were vernalized 96hrs and grown with 16/8 hrs light/dark at 21°C for 7 days and the seedlings were moved to MS or MS+IAA plates and grown vertically for 7 days.

### Cloning and transgenic lines

*PATs* promoter sequences with 5’ HindIII and 3’ XbaI linkers were amplified from Col-0 genomic DNA and cloned in plasmid pGWB515 to make pGWB515-PATsprom [61]. The Venus sequence without stop codon was amplified from pEN-L1-Venus-L2 [62] and cloned in pGWB515-PATsprom. PAT genes were amplified from Col-0 genomic DNA and cloned into pENTR-D-TOPO (Invitrogen). The entry clones were combined with pGWB515-PATsprom and pGWB515-PATsprom-Venus to obtain N-terminal HA and Venus tags respectively [61]. pENTR-D-TOPO-GUS (Invitrogen) and pENTR-D-TOPO-LSM1 [26] were combined to pK7WGY2.0 [63] to obtain N-terminal YFP tags. The PATH1 promoter was also cloned in pK7WGC2.0 to obtain pK7WGC2.0-PATH1prom, then combined with pENTR-D-TOPO-PATH1 to produce PATH1 with an N-terminal CFP tag. These fusions were transformed into *Agrobacterium tumefaciens* strain GV3101 for transient and stable expression. Arabidopsis transformation was by floral dipping [64]. *arr10/arr12/oxASL9* was generated by vacuum infiltrating *arr10/arr12* with *A. tumefaciens* strain EHA101 harbouring pSK1-ASL9[44]. Transformants were selected on hygromycin (30 mg/L, for pGWB515) or kanamycin (50mg/L, for pK7WGC2.0) MS agar, and survivors tested for transcript expression by qRT-PCR and protein expression by immuno-blotting. Primers for cloning are provided in Table S1

### Transient expression, protein extraction and co-IP in *Nicotiana benthamiana*

*A. tumefaciens* strains carrying PAT fusions were grown in YEP medium supplemented with appropriate antibiotics overnight. Cultures were centrifuged and re-suspended in buffer (10mM MgCl2, 10 mM MES-K (pH 5.6), 100 μM acetosyringone) to OD600=0.8. *A. tumefaciens* strains carrying PATs-HA and YFP-LSM1 or YFP-GUS were mixed 1:1 and infiltrated into 3 week-old *N. benthamiana* leaves. Leaf samples for protein extraction and immunoprecipitation were collected 3 days post infiltration (dpi). Tissues for protein extraction were ground in liquid nitrogen and IP buffer (50mM Tris-HCl pH 7.5; 150 mM NaCl; 5 % (v/v) glycerol; 1 mM EDTA; 0.1%(v/v) NP40; 10 mM DTT; protease inhibitor cocktail (Roche); Phosstop (Roche)) added at 2mL/g tissue powder. Following 20 min centrifugation at 4°C and 13000 rpm, sample supernatants were adjusted to 2mg/ml protein and incubated 4 hours at 4°C with GFPTrap-A beads (Chromotek). Beads were washed 4 times with wash buffer (20 mM Tris pH 7.5; 150m M NaCl; 0.1 %(v/v) NP40 before adding 4×SDS buffer (novex)) and denatured by heating at 95°C for 5 min.

### Protein extraction, SDS-PAGE and immunoblotting

Tissue was ground in liquid nitrogen and 4×SDS buffer (novex) was added and heated at 95°C for 5 min, cooled to room temperature for 10min, samples were centrifuged 5min at 13000 rpm. Supernatants were separated on 10% SDS-PAGE gels, electroblotted to PVDF membrane (GE Healthcare), blocked in 5% (w/v) milk in TBS-Tween 20 (0.1%, v/v) and incubated 1hr to overnight with primary antibodies (anti-GFP (AMS Biotechnology 1:5.000, anti-HA 1:1,000 (Santa Cruz)). Membranes were washed 3 × 10 min in TBS-T (0.1%) before 1hr incubation in secondary antibodies (anti-rabbit or anti-mouse HRP or AP conjugate (Promega; 1: 5.000)). Chemiluminescent substrate (ECL Plus, Pierce) was applied before camera detection. For AP-conjugated primary antibodies, membranes were incubated in NBT/BCIP (Roche) until bands were visible.

### Confocal microscopy

Imaging was done using a Leica SP5 inverted microscope. The confocal images were analysed with Zen2012 (Zeiss) and ImageJ software. Representative maximum intensity projection images of 10 Z-stacks per image have been shown in Fig2.

### RNA extraction and qRT-PCR

Total RNA from tissues was extracted with TRIzol^™^ Reagent (Thermo Scientific), 2μg total RNA were treated with DNAse I (Thermo Scientific) and reverse transcribed into cDNA using RevertAid First Strand cDNA Synthesis Kit according to the manufacturer’s instructions (Thermo Scientific). The *ACT2* gene was used as an internal control. qPCR analysis was performed on a Bio-RAD CFX96 system with SYBR Green master mix (Thermo Scientific). Primers are listed in Table S1. All experiments were repeated at least three times each in technical triplicates.

### XRN1 assay

Total RNA was extracted from tissues using the NucleoSpin^®^ RNA Plant kit (Machery-Nagel). 1μg RNA was incubated with 1 unit XRN1 (New England Biolabs) or water for 2hr at 37°C. RNA was then reverse transcribed into cDNA with RevertAid First Strand cDNA Synthesis Kit (Thermo Scientific). Target transcript accumulation was measured by qPCR using SYBR Green master mix (Thermo Scientific) and normalized to *ACT2*. Calculating 5’ capped versus total transcripts was done by comparing transcript levels from XRN1 and mock-treated samples for the individual genotypes [26, 65].

### RIP assay

RIP was performed as previously described [66]. 1.5g tissues were fixated by vacuum infiltration with 1% formaldehyde for 20min followed by 125 mM glycine for 5min. Tissues were ground in liquid nitrogen and RIP lysis buffer (50mM Tris-HCl pH 7.5; 150mM NaCl; 4mM MgCl2; 0.1% Igepal; 5 mM DTT; 100 U/mL Ribolock (Thermo Scientific); 1 mM PMSF; Protease Inhibitor cocktail (Roche)) was added at 1.5mL/g tissue powder. Following 15 min centrifugation at 4°C and 13000rpm, supernatants were incubated with GFPTrap-A beads (Chromotek) for 4 hours at 4°C. Beads were washed 3 times with buffer (50 mM Tris-HCl pH 7.5; 500 mM NaCl; 4 mM MgCl2; 0.5 % Igepal; 0.5 % Sodium deoxycholate; 0.1 % SDS; 2 M urea; 2 mM DTT before RNA extraction with TRIzol^™^ Reagent (Thermo Scientific)). Transcript levels in input and IP samples were quantified by qPCR, and levels in IP samples were corrected with their own input values and then normalized to YFP WAVE lines for enrichment.

### RNA-seq analysis

Total RNA was extracted from 6 week-old soil grown plants using the NucleoSpin^®^ RNA Plant kit (Machery-Nagel). RNA quality, library preparation and sequencing were performed by BGI. RNA-seq reads were mapped to the *Arabidopsis thaliana* TAIR10 reference genome with STAR (version 2.5.1b) using 2-pass alignment mode[67]. The read counts for each gene were detected using HTSeq (version 0.5.4p3)[68]. The Araport11 annotation was used for both mapping and read counting. The counts were normalized using the TMM normalization from edgeR package in R. Prior to statistical testing the data was voom transformed and then the differential expression between the sample groups was calculated with limma package in R. Genes with fold change ≥2 or ≤-2 and P-value ≤0.01 are listed in Data File S1. Functional Annotation Tool DAVID Bioinformatics Resources 6.8 has been used for GO term analysis [37].

### Statistical analysis

Statistical details of experiments are reported in the figures and legends. Systat software was used for data analysis. Statistical significance between groups was determined by one-or two-way ANOVA (analysis of variance) followed by Holm-Sidak test.

## Supporting information

Data Set S1

---

**Fig S1.**
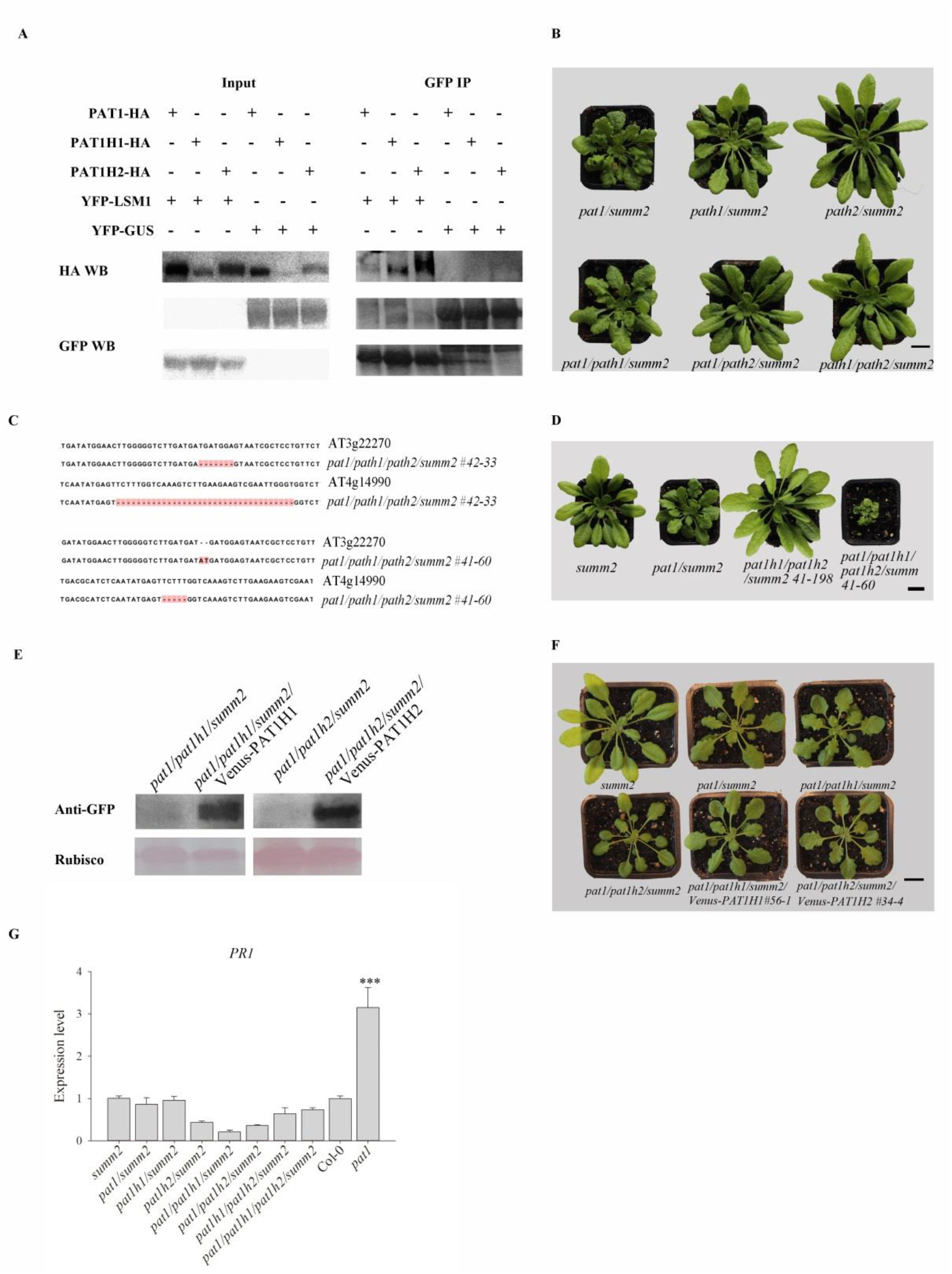
Characterization of *pat* mutants generated using CRISPR/CAS9 system. (**A**), Co-IP between the three PAT-HA and YFP-LSM1 fusions. Proteins were transiently co-expressed in *N*. *benthamiana* and tissue harvested 3 days post-infiltration. Immunoblots of inputs (left panels) and GFP IPs (right panels) were probed with anti-HA antibodies and anti-GFP antibodies. (**B**), 6 week-old plants of *pat1/summ2*, *path1/summ2, path2/summ2, pat1/path1/summ2, pat1/path2/summ2* and *path1/path2/summ2* grown in soil in a chamber with 8/16hrs light/dark at 21°C. One representative plant for each line is shown. The scale bar indicates 1cm.(**C**), Sequencing of *path1* and *path2* mutations in independent lines *pat1/path1/path2/summ2* 42-33 and *pat1/path1/path2/summ2 41-60*. (**D**), 6 week-old plants of *summ2*, *pat1/summ2*, *path1/path2/summ2 41-198* and *pat1/path1/path2/summ2 41-60* grown in soil in chamber with 8/16hr light/dark at 21°C. One representative plant for each line is shown. The scale bar indicates 1cm. Western blots detecting the expression of PATH1 and PATH2 fusions with N-Venus (**E**) and growth phenotype (**F**) of complemented lines. One representative plant for each line is shown. The scale bar indicates 1cm.(G) PR1 expression level in 5 week-old plants of *pats* mutant in summ2 background, *pat1* and Col-0, the experiment was repeated 3 times, bars marked with three asterisk (***) are statistically extremly significant (P-value <0.001).

**Fig S2.**
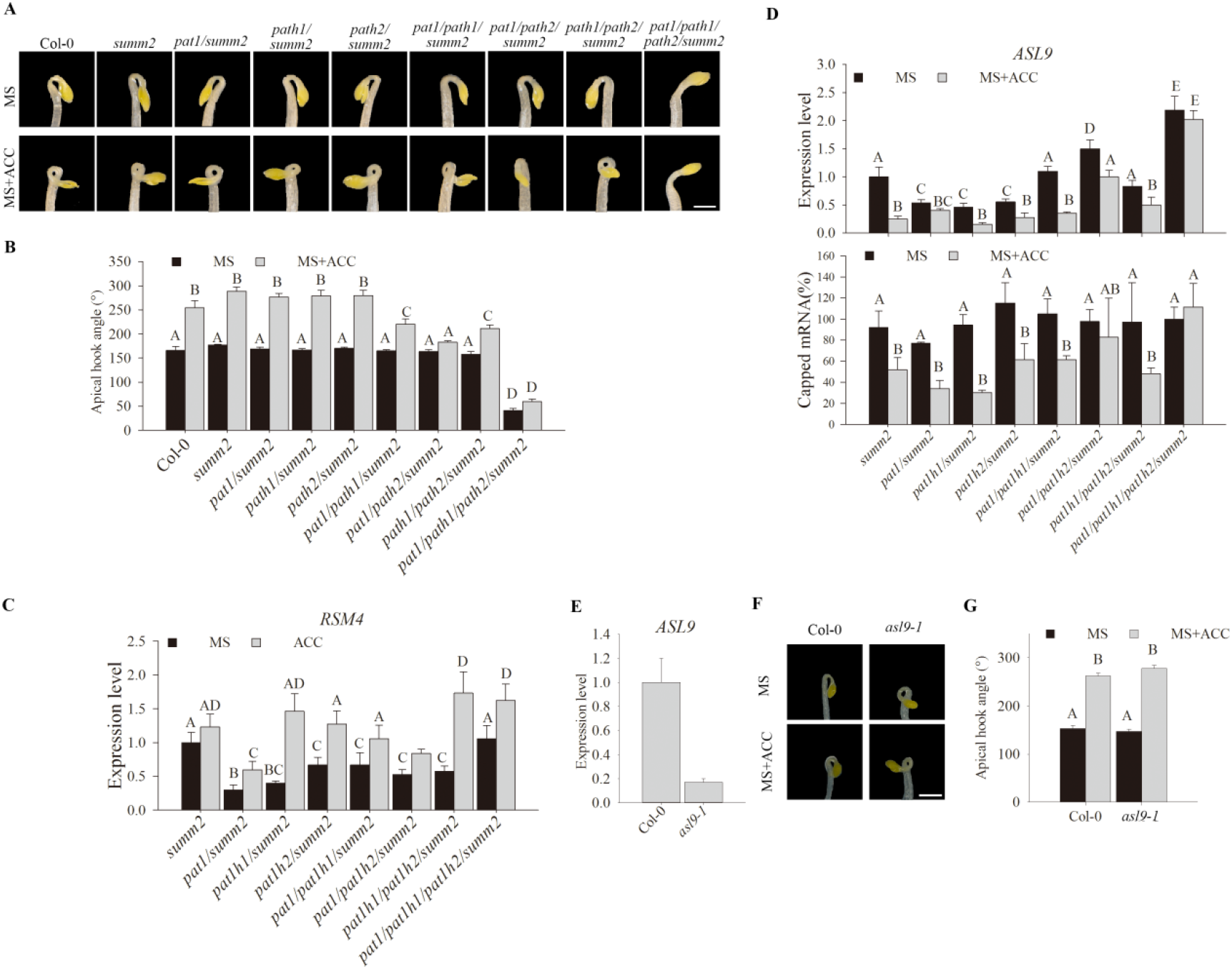
Apical hook in *pat(s)* and *asl9-1* mutants. Hook phenotypes (**A**) and apical hook angles (**B**) in triple responses to ACC treatment of etiolated seedlings of Col-0, *summ2*, *pat1/summ2, path1/summ2, path2/summ2, pat1/path1/summ2, pat1/path2/summ2, path1/path2/summ2* and *path1/path2/summ2*. The treatment was repeated 3 times, and representative pictures are shown. The scale bar indicates 1mm. (**C**) *RSM4* expression levels in cotyledons and hook regions of dark-grown *pat* single, double and triple mutants under control or ACC treatment. Error bars indicate SE (n = 3). One representative of 3 biology replicates is shown. (**D**) *ASL9* expression levels and capped *ASL9* transcript levels in cotyledons and hook regions of dark-grown *pat* single and double mutants under control or ACC treatment. Experiments were performed together with Fig 4A. Error bars indicate SE (n = 3). One representative of 3 biology replicates is shown. (**E**) *ASL9* expression levels in 4 week-old plants of Col-0 and *asl9-1*. Experiments were performed together with Fig 4A. Error bars indicate SE (n = 3). One representative of 3 biology replicates is shown. Hook phenotypes (**F**) and apical hook angles (**G**) in triple responses to ACC treatment of etiolated Col-0 and *asl9-1* seedlings. The treatment was repeated 3 times, and representative pictures are shown. The scale bar indicates 1mm.

**Fig S3.**
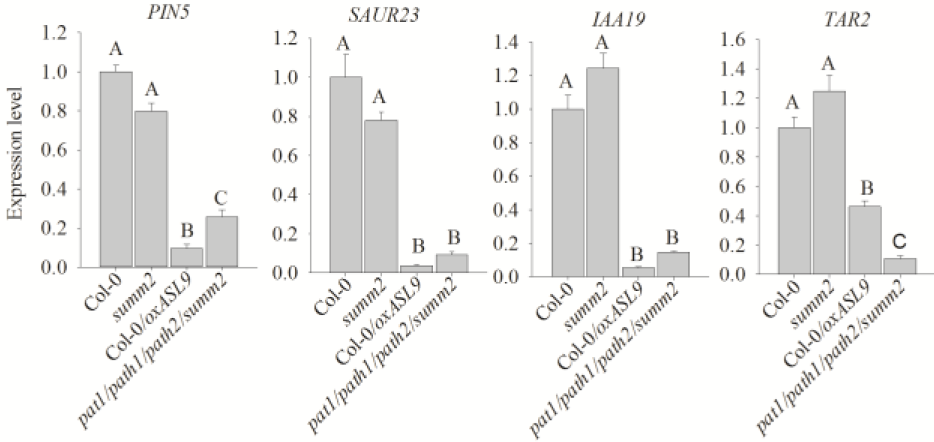
Auxin related genes expression in mRNA decay deficient mutant and *ASL9* over-expressor. Auxin pathway genes *PIN5, SAUR23, IAA19 and TAR2* expression levels in 10 day-old seedlings of Col-0, *summ2*, Col-0/*oxASL9* and *pat1/path1/path2/summ2*. The experiment was repeated 3 times, and representative pictures are shown. Bars marked the same letter are not significantly different from each other (P-value>0.05).

**Fig S4.**
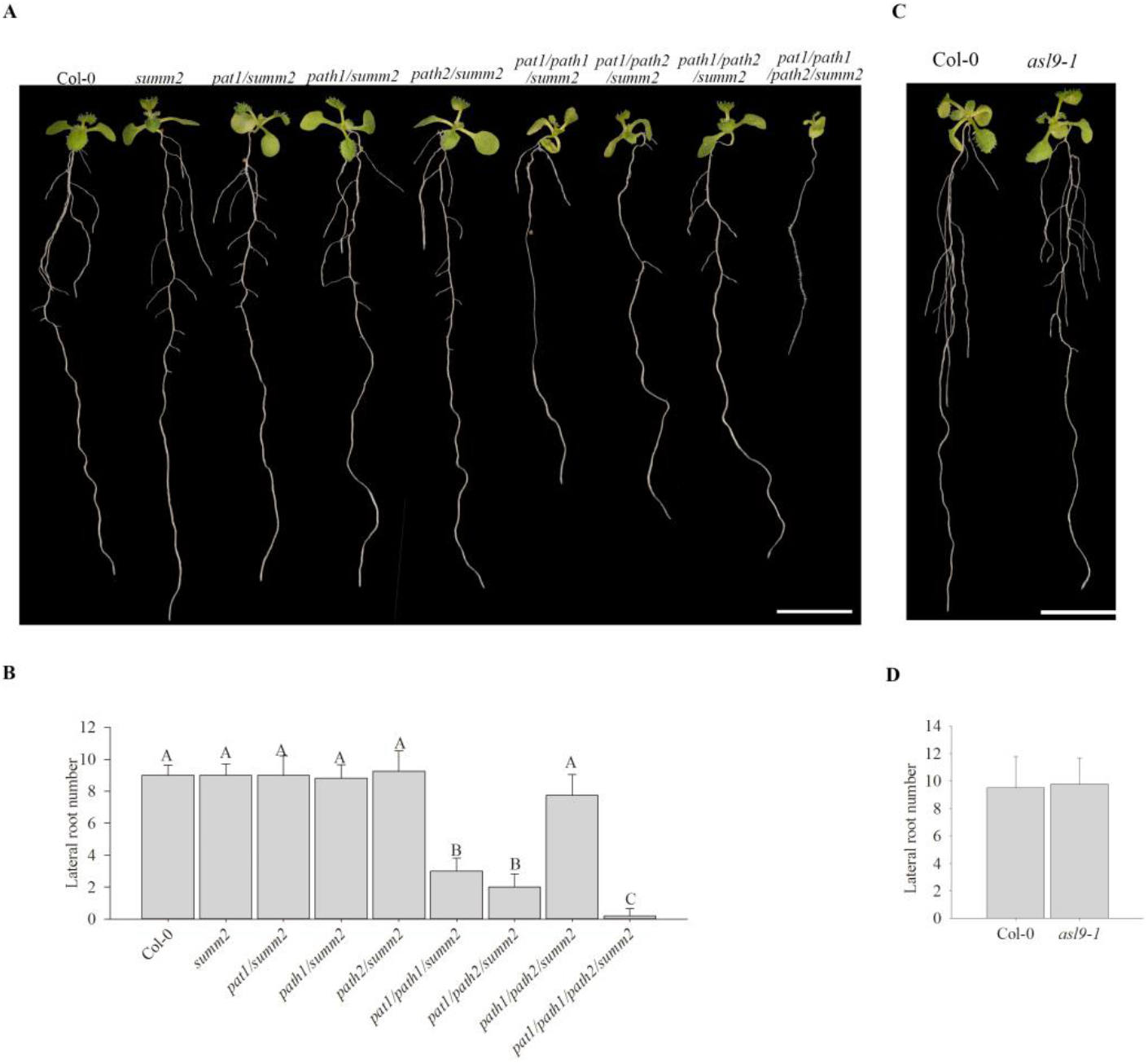
LR formation in *pat(s)* and *asl9-1* mutants. Phenotypes (**A**) and LR number counts (**B**) of 10-day old seedlings of *pat1/summ2*, *path1/summ2, path2/summ2, pat1/path1/summ2, pat1/path2/summ2* and *path1/path2/summ2*. Phenotypes (**C**) and LR number counts (**D**) of 10-day old seedlings of Col-0 and *asl9-1*. The treatment was repeated 4 times, and representative pictures are shown. The scale bar indicates 1cm. Bars marked with asterisks are statistically significant (**:P-value <0.01; ***: P-value<0.001).

**Fig S5.**
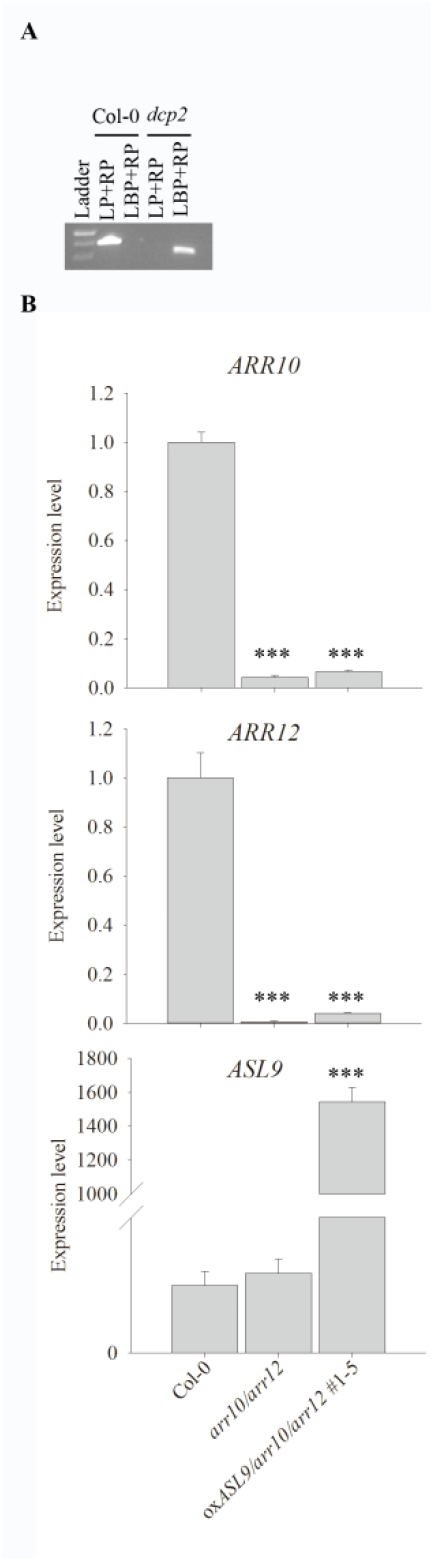
Characterization of *dcp2* and *arr10/arr12/oxASL9* mutants. (**A**) Determining the genotype of hookless seedlings germinated from *dcp2* heterozygote seeds. PCR was carried out on genomic DNA from Col-0 and hookless seedlings germinated from *dcp2* heterozygote seeds using primer DCP2LP and DCP2RP to detect plants that contained a wild-type copy of *DCP2*, LBb1.3 for the T-DNA and gene-specific primer DCP2R were used to detect the presence of T-DNA. Annotation in this analysis is indicated: LP, DCP2LP; RP, DCP2RP; LBP, LBb1.3. (**B**) *ARR10*, *ARR12* and *ASL9* expression levels in Col-0, *arr10/arr12* and *arr10/arr12/oxASL9* 1-5 seedlings. The experiment was repeated 3 times, and representative pictures are shown. Bars marked with three asterisk (***) are statistically extremly significant (P-value <0.001).

**Fig S6.**
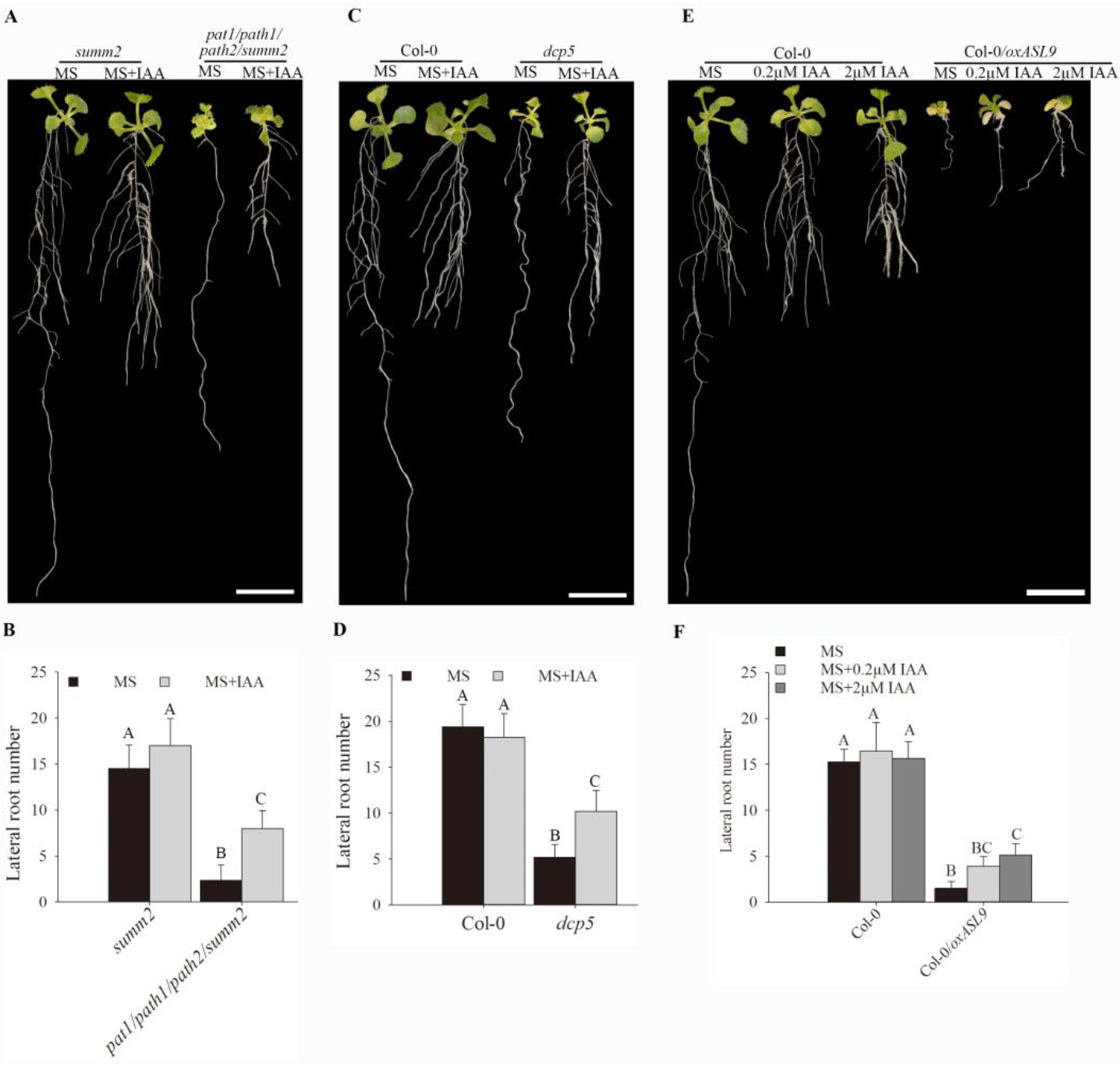
Auxin restores LR formation in mRNA decay deficient mutants and Col-0/*oxASL9*. Phenotypes(**A**) and LR number counts(**B**) of 14-day old seedlings of *summ2* and *pat1/path1/path2/summ2* on MS or MS with 0.2μM IAA. Phenotypes (**C**) and LR number counts (**D**) of 14-day old seedlings of Col-0 and *dcp5* on MS or MS with 0.2μM IAA. Phenotypes (**E**) and LR number counts (**F**) of 14-day old seedlings of Col-0 and Col-0/*oxASL9* on MS, MS with 0.2μM IAA or MS with 2μM IAA. Seeds on MS plates were vernalized 96hrs and grown with 16/8 hrs light/dark at 21°C for 7 days. The seedlings were moved to MS or MS+IAA plates and grown vertically for 7 days. The treatment was repeated 3 times, and representative pictures are shown. The scale bar indicates 1cm. Bars marked the same letter are not significantly different from each other (P-value>0.05).

## Acknowledgments

We thank Qi-Jun Chen for Phee401 and Nam-Hai Chua for *dcp5* and *dcp2* seeds. Special thanks to John Mundy for advice throughout the project and critically reading the manuscript. We acknowledge the Bioinformatics and Scientific Computing Facility (VBCF) for the next-generation sequencing data analysis. This work was supported by the Novo Nordisk Fonden to MP (NNF18OC0052967), the Institute Strategic Programme grant (BB/P013511/1) to the John Innes Centre and a PhD scholarship from China Scholarship Council to ZZ (201504910714).

ZZ, MER, and MP conceived and designed the experiments. ZZ, MER, JRC, YD, TY, SDH, EK and LØ performed experiments. ZZ and MP analyzed the data. ZZ and MP wrote the manuscript.

The authors declare no competing interests.

Correspondence and requests for materials should be addressed to MP.

